# Effects of catechol-*O*-methyltransferase inhibition on effort-related choice behavior in male mice

**DOI:** 10.1101/2020.01.28.924142

**Authors:** Adrienne DeBrosse, Abigail M. Wheeler, James C. Barrow, Gregory V. Carr

## Abstract

Effort-related choice (ERC) tasks allow animals to choose between high-value reinforcers that require high effort to obtain or low-value/low-effort reinforcers. Dopaminergic neuromodulation regulates effort-related choice behavior. The enzyme catechol-*O*-methyltransferase (COMT) metabolizes synaptically-released dopamine. COMT is the predominate regulator of dopamine turnover in regions of the brain with low levels of dopamine transporters, including the prefrontal cortex. Here, we evaluated the effects of the COMT inhibitor tolcapone on ERC performance in a touchscreen-based fixed-ratio/concurrent chow assay in male mice. In this task, mice were given the choice between engaging in a fixed number of instrumental responses to acquire a strawberry milk reward and consuming standard lab chow concurrently available on the chamber floor. We found no significant effects of tolcapone treatment on either strawberry milk earned or chow consumed compared to vehicle treatment. In contrast, we found that haloperidol decreased instrumental responding for strawberry milk and increased chow consumption as seen in previously published studies. These data suggest that COMT inhibition does not significantly affect effort-related decision making in a fixed-ratio/concurrent chow task in male mice.

## Introduction

Humans and other mammals interact with complex stimuli in their environments and make decisions to maximize economic value and improve survival odds. Economic value is defined as the benefit provided by a good or service relative to the amount of currency required to procure it (Samuelson, 1947). A common type of decision is one where a high-value item and a low-value item of a similar quality are both available. The choice is easy when the amount of effort (currency) required to procure both items is the same. Conversely, when the high-value option requires more work to attain, the calculation becomes more complicated because effort-based discounting of the high-value option determines the choice (Botvinick et al 2009, Walton et al 2006).

The capacity to make effort-based decisions is disrupted in many psychiatric and neurological disorders, including depression and schizophrenia (Barch et al 2014, Fervaha et al 2013, Gold et al 2013, Treadway et al 2012). In the case of schizophrenia, the cognitive effort allocated to gaining a reward drops off more steeply than in healthy individuals, and is correlated with overall negative symptomatology (Culbreth et al 2016). These data suggest that increased understanding of how effort-based decisions are made would potentially improve treatment options for psychiatric disorders.

There are several paradigms for assessing effort-based decision-making in humans and animal models, including the human Effort Expenditure for Rewards Task (EEfRT), rodent T-maze with barriers, and rodent operant concurrent choice tasks (Botvinick et al 2009, Cousins et al 1994, Enomoto et al 2018, Mott et al 2009, Treadway et al 2012). Recently, rodent touchscreen-based versions of the operant concurrent choice effort-related choice (ERC) tasks have been developed (Heath et al 2015, Yang et al 2020).

The dopaminergic system has long been recognized as a critical regulator of ERC performance from studies that utilized systemic manipulations. The D2 antagonist haloperidol biases behavioral toward the low-effort low value option across multiple versions of ERC tasks (Randall et al 2012, Salamone et al 1994, Salamone et al 1991, Yang et al 2020). Similarly, the D1 receptor antagonist ecopipam decreases choice of the high-effort arm of a T-maze and decreases lever pressing in a fixed ratio (FR)/concurrent chow paradigm (Yohn et al 2015a). Modulation of dopamine transporter (DAT) activity also affects ERC performance. DAT knockdown mice demonstrate increase preference for the high-effort high-value reward in an FR/concurrent chow task (Cagniard et al 2006). Pharmacological treatment with DAT inhibitors also biases rats toward the high-effort reward in a progressive ratio (PR)/concurrent chow task (Sommer et al 2014).

Experiments targeting the nucleus accumbens have demonstrated a particularly strong involvement of the mesolimbic dopaminergic system in ERC behavior. Dopamine depletion in the nucleus accumbens (NAc) biases responding toward the low-effort low-reward option (Cousins et al 1996, Salamone et al 1994). Local injections of both the D1 antagonist SCH 23390 and the D2 antagonist raclopride into the NAc shell and core decrease lever pressing in rats during a FR/concurrent chow task (Nowend et al 2001). Direct injections of haloperidol into the NAc likewise decrease lever pressing on a FR5/concurrent chow task (Salamone et al 1991).

The relationship between NAc dopamine and ERC performance is bidirectional as increased dopaminergic signaling biases responding toward the high-effort high-reward options. Overexpression of the D2 receptor in the NAc increases lever pressing on progressive (PR) and an ERC task (Trifilieff et al 2013). Individual rats that respond more highly at baseline during a progressive ratio/concurrent chow lever pressing task show higher levels of D1 receptor activation, measured as changes in DARPP-32 phosphorylation, in NAc core (Randall et al 2012). Moreover, in human fMRI studies, the NAc was activated during an effort discounting task in direct relation to the amount of reward that was cued (Botvinick et al 2009).

The role of DA in the prefrontal cortex in ERC performance is less clear. Rodent lesion and human fMRI studies implicate the anterior cingulate cortex (ACC) in regulating ERC performance. Specifically, ACC lesions bias responding toward the low-effort low-value options (Walton et al 2003, Walton et al 2002) and the ACC is activated in fMRI studies during effort-based decision-making (Botvinick et al 2009, Klein-Flugge et al 2016). The D1 antagonist SCH23390 reduced preference for the high-effort high value reward (Schweimer & Hauber 2006). However, the effects of catecholamine depletion in the ACC are ambiguous as one study found no effect (Walton et al 2005) while a separate study found that depletion reduced preference for the high-effort high-value option (Schweimer et al 2005).

To our knowledge, there have not been any studies investigating the effects of augmenting PFC dopaminergic activity on ERC performance. The enzyme catechol-*O*-methyltransferase (COMT) is an important regulator of cortical dopaminergic function due to its role in the degradation of synaptically-released dopamine (Mannisto & Kaakkola 1999). Inhibition of COMT leads to a decrease in dopamine turnover in regions of the brain with low DAT expression, including the PFC (Kaenmaki et al 2010, Yavich et al 2007). COMT also plays a significant role in intertemporal choice and delay discounting (Boettiger et al 2007, Kayser et al 2012), but it has not been investigated in an ERC task. Here, we tested the effects of the COMT inhibitor tolcapone in a touchscreen-based ERC task.

## Materials and Methods

### Mice

Eight-week-old male C57BL/6J mice (The Jackson Laboratory, Bar Harbor, ME, USA) were used in all experiments. The mice were housed in disposable polycarbonate caging (Innovive, San Diego, CA, USA) in groups of four. The mice were maintained on a 12/12 light/dark cycle (lights on at 0600 hours). Water was available in the home cage *ad libitum* throughout all experiments. The mice were fed Teklad Irradiated Global 16% Protein Rodent Diet (#2916; Envigo, Indianapolis, IN, USA) in the home cage *ad libitum* until the start of the food restriction protocol. Two separate cohorts of mice were tested in these experiments and all testing was done during the light phase (1200-1600 hours).

### Drugs

Tolcapone was synthesized in-house. It was suspended in vehicle (0.1% Tween80, 0.1% 1510 silicone antifoam, 1% methylcellulose in water). Haloperidol was purchased from Sigma-Aldrich (St. Louis, MO, USA). It was dissolved in 1% DMSO in a citrate phosphate buffer (pH 5). Both drugs were administered by intraperitoneal (ip) injection one hour before the start of behavioral testing. Administration volumes were 10 mL/kg.

### Food Restriction Protocol

Upon arrival at the animal facility, mice were given at least 72 hours to acclimate to the colony room before handling by experimenters. Mice were handled and weighed daily from that point forward. After at least two days of handling, mice were food restricted to 3g of chow per mouse per day in order to maintain 85-90% of their predicted free-feeding weight based on average growth curve data (The Jackson Laboratory). Mice were removed from the standard group housing setup due to aggression or significant deviations from the expected body weight during food restriction. In Cohort A, we split one mouse that was under 85% of the predicted average body weight and two mice that were >90% predicted average body weight while their cage mates were in range. In Cohort B, two mice were split and single housed. To familiarize the mice with the Nesquik strawberry milk (Nestlé, Vevey, Switzerland) reward used in the effort-related choice task, we introduced the milk to the home cage on 4×4 inch weighing paper (VWR, Radnor, PA, USA). The weighing paper was left in the cage until all mice had sampled the strawberry milk. This procedure was repeated for a total of two days.

### Operant Chamber Habituation and Initial Touch Training

We adapted the effort-related choice (ERC) training and task parameters described by Heath and colleagues (Heath et al 2015). Our training and test schedules were programmed in the Animal Behavior Environment Test System (ABET II; Lafayette Instrument, Lafayette, IN, USA) and run in four Bussey-Saksida Mouse Touch Screen Chambers (Model 80614E, Lafayette Instrument, Lafayette, IN, USA). All training and testing was conducted five days per week (Monday-Friday). Mice were habituated to the chambers during two consecutive 20-minute sessions in which they were presented with 200 μL of strawberry milk in the reward tray. Mice passed the Habituation stage when they had consumed all of the milk in the tray. After reaching criterion, mice advanced to the Initial Touch stage. During Initial Touch training, mice could receive reward by making a nose poke response at the center square in a 5-mask array. Nose pokes resulted in receiving 30 μL of strawberry milk while omissions resulted in 10 μL of reward at the end of the 30-second trial timer. For Initial Touch training, passing criterion was set at 30 completed trials within the 60-minute session.

### Fixed Ratio Training

During Fixed Ratio (FR) training, mice were required to nose poke the illuminated center square of the 5-mask array in order to receive reward (10 μL). Each session had an assigned response requirement, or number of touches to the center square that were needed to earn each reward. For example, during an FR2 schedule a mouse would receive one reward after responses at the center square. Response requirement remained constant within a session and increased as mice passed through training. We trained the mice on response requirements of one (FR1), two (FR2), three (FR3), and five (FR5) responses. Criterion was set at 30 completed trials within a 30-minute session. Mice were required to reach criterion on one FR1, one FR2, one FR3, and three consecutive FR5 sessions in order to advance on to the effort-related choice task. In cases where mice reached the FR5 criterion on days other than Friday, we maintained mice on the FR5 schedule until Monday of the following week in order avoid confounding schedule and day of the week in the next phase of the experiment.

### The Effort-Related Choice Task

The effort-related choice (ERC) task utilized FR schedules with the addition of a preweighed ~3 g pellet of standard laboratory chow to the floor of the touchscreen chamber. Before the mouse was loaded into the chamber, chow was placed between the front and back infrared beams. In order to counterbalance starting chow position across chambers, chow was placed next to either the back or front wall on the right or left side of the chamber. Chow location was counterbalanced across mice. Mice had the choice of responding at the touchscreen in order to earn strawberry milk rewards based on the specific FR schedule or consume the freely available chow. Each session was 30 minutes in duration. At the conclusion of each session, chow crumbs were swept down into the catch tray beneath the chamber and weighed together with remaining chow to determine how much was consumed during the session.

The mice in these experiments were split into two cohorts that were tested on separate protocols to answer specific experimental questions.

### Cohort A

During the first phase of ERC testing, the mice were tested on FR schedules with increasing response requirements over five days (FR1, FR2, FR4, FR8, and FR16). The same schedules were presented during the next week but in reverse order (FR16, FR8, FR4, FR2, FR1). Next, we tested the effect of tolcapone on choice at three doses (3, 10, and 30 mg/kg) using the FR4 schedule over the following three weeks. We interleaved drug/vehicle dosing days with no injection baseline days. We also tested haloperidol (0.1 mg/kg) as a comparator to validate our protocol.

### Cohort B

This cohort proceeded directly from training to the drug treatment phase. The drug treatment phase was identical to Cohort A.

### Statistical Analysis

All statistical analyses were conducted in GraphPad Prism 8 (GraphPad Software, San Diego, CA, USA). We used t-tests to analyze the habituation and training data. Welch’s t-tests were used for the Habituation, Initial Operant, FR2, and FR3 stages due to unequal variances between the groups. The baseline ERC and ERC-tolcapone data analyzed using repeated measures one-way ANOVAs with Geisser-Greenhouse corrections. We utilized paired t-tests to analyze the haloperidol data. Tukey’s tests were used for post hoc analyses. In analyses where performance following drug or vehicle treatment was compared to baseline performance, the baseline was calculated by averaging performance from the three baseline testing days during the same week. One mouse was removed from the analyses of the tolcapone experiment because it did not receive the 30 mg/kg dose due to experimenter error. The statistical significance threshold was set at p < 0.05. Data are presented as the mean ± SEM.

## Results

### Training

All mice progressed through training and advanced to the ERC task. Data for the training stages are shown in Table 1. There was no difference in the number of sessions to reach criterion during the Habituation stage (*t_7_* = 1.871, *p* = 0.1036), however, Cohort B required more total sessions to pass the Initial Operant (*t_7_* = 3.416, *p* = 0.0112) and the FR1 (*t_14_* = 3.326, *p* = 0.0050) stages. These group differences did not persist beyond these early training stages as there were no differences in the sessions required to reach criterion in the FR2 (*t_7_* = 1.000, *p* = 0.3506), FR3 (*t_7_* = 2.049, *p* = 0.0796), or FR5 (*t_14_* = 0.8215, *p* = 0.4251) stages.

**Table 1.** Days to Criterion

| Cohort | Habituation | Initial Operant | FR1 | FR2 | FR3 |
| --- | --- | --- | --- | --- | --- |
| A | 2.0 ± 0 | 1.0 ± 0 | 2.2 ± 0.25 | 1.2 ± 0.25 | 1.0 ± 0 |
| B | 2.5 ± 0.27 | 1.6 ± 0.18* | 4.0 ± 0.46** | 1.0 ± 0 | 1.8 ± 0.37 |

\* $p < 0.05$ and \*\* $p < 0.01$ compared to Cohort A
n = 8/cohort

### Baseline ERC performance

Cohort A was first tested on an ERC task where the ratio requirement to earn the strawberry milk reward was varied across sessions. Mice had two sessions each with four different fixed ratio requirements (FR1, FR2, FR4, FR8, and FR16). For analysis, we averaged the two sessions for each mouse. As expected, the number of rewards earned, expressed as strawberry milk consumed, decreased as the ratio requirement increased (*F_4, 28_* = 108.7, *p* < 0.0001). The mean value of milk consumed at each ratio requirement was significantly different from all of the other ratios tested (Figure 1). Also, as the work requirement for the strawberry milk increased, consumption of the freely available chow increased (*F_4, 28_* = 4.380, *p* = 0.0071; Figure 1). Post hoc comparisons showed that the amount of chow consumed during FR1 sessions was significantly lower than the amount consumed on FR8 (*p* =0.0465) and FR16 (*p* =0.0047) sessions.

**Figure 1.**
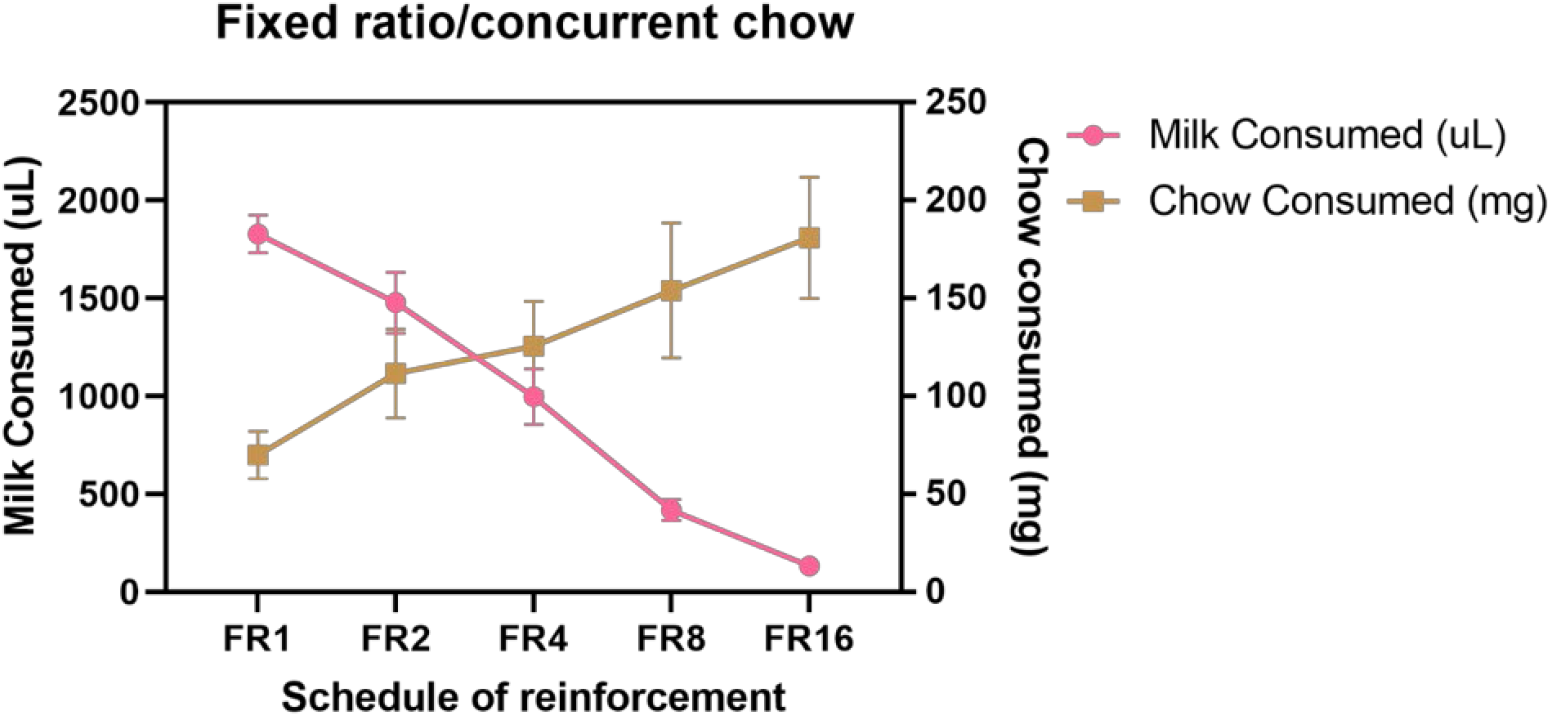
Baseline performance on variable FR/concurrent chow sessions. Strawberry milk consumed is graphed on the left axis and chow consumed is graphed on the right axis. Each mouse was tested twice at each FR requirement. The two sessions were averaged to produce a single value for each mouse. Strawberry milk consumption under each FR requirement was significantly different from all of the others. All comparisons were *p* < 0.0001 except for FR1 vs. FR2 (*p* = 0.0087) and FR8 vs. FR16 (*p* = 0.0426). Chow consumption increased as the FR requirement for strawberry milk increased. Chow consumption in the FR8 (*p* = 0.0465) and FR16 (*p* = 0.0047) sessions were significantly different then the FR1 session. \**p* < 0.05; \*\**p* = 0.01. N = 8.

### Effects of tolcapone on ERC performance

For the drug treatment stage of testing, we chose to use the FR4 schedule for all sessions. Based on the data collected in the first experiment, the FR4 schedule was predicted to produce a baseline level of performance that would allow us to measure significant increases or decreases in both milk and chow consumption.

Tolcapone did not alter the amount of strawberry milk earned (*F_1.838, 25.74_* = 1.153, *p* = 0.3272) or chow consumed (*F_2.294, 32.12_* = 2.331, *p* = 0.1068) at any dose tested (3, 10, 30 mg/kg; Figure 2a-b). There were significant differences between individual mice in both strawberry milk earned (*F_14, 42_* = 10.26, *p* < 0.0001) and chow consumed (*F_14, 42_* = 2.864, *p* = 0.0042). Because of these individual differences and fluctuations in baseline performance across the weeks of testing, we also analyzed the tolcapone data based on % change from each mouse’s baseline strawberry milk and chow consumption. This reanalysis did not change the results of the comparison. There were still no significant changes in strawberry milk earned (*F_2.247, 31.45_* = 1.621, *p* = 0.2118; Figure 2c) or chow consumed (*F_1.044, 14.62_* = 0.7226, *p* = 0.4149; Figure 2d).

**Figure 2.**
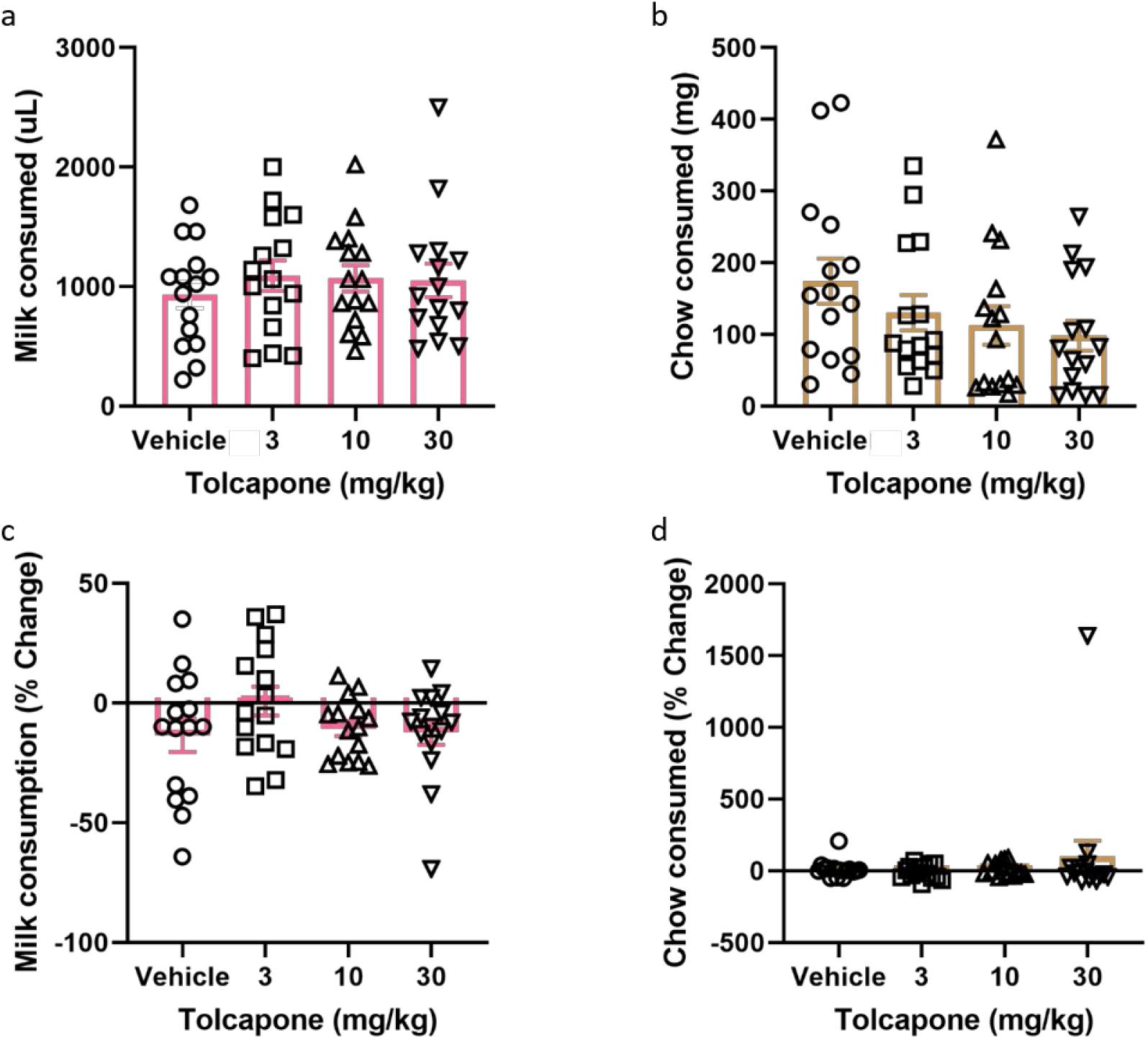
Effects of tolcapone on performance in FR4/concurrent chow sessions. Tolcapone (3, 10, and 30 mg/kg) did not affect strawberry milk (a) or chow consumed (b). We also analyzed the data by normalizing performance to baseline (no vehicle or drug injection) sessions. After normalization, there was still no effect of tolcapone on strawberry milk (c) or chow consumed (d). N = 15.

### Effects of haloperidol on ERC performance

In order to validate our ERC protocol, we also tested the effects of haloperidol (0.1 mg/kg) as a positive control. Haloperidol has been shown to decrease responding for the high value option in ERC, including a recent study using a touchscreen-based task (Yang et al 2020). Here, we found that haloperidol decreased the amount of strawberry milk earned (*t_15_* = 2.477, *p* = 0.0272; Figure 3a), but did not significantly increase the amount of chow consumed (*t_15_* = 1.685, *p* = 0.1126; Figure 3b).

**Figure 3.**
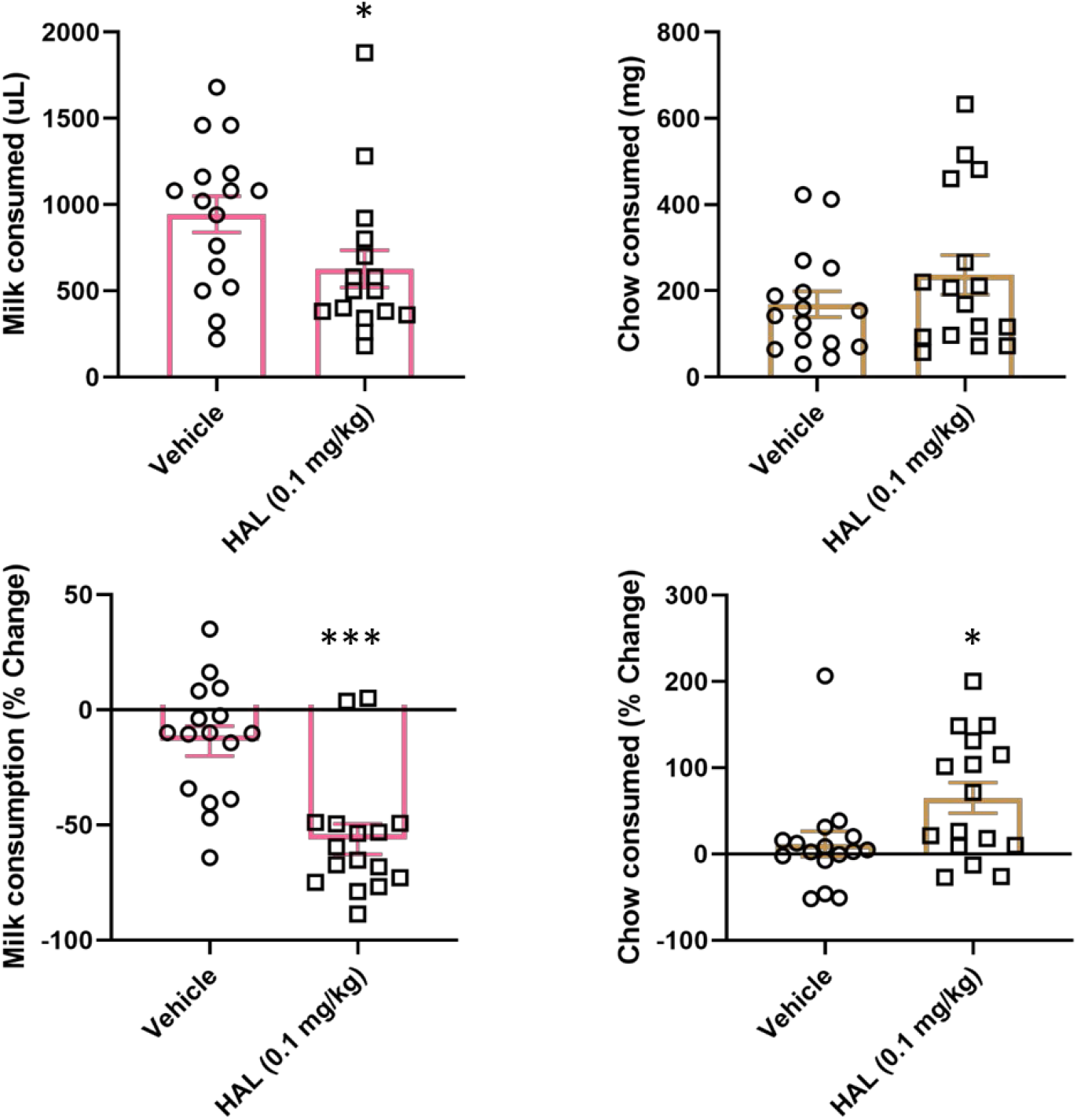
Effects of haloperidol on performance in FR4/concurrent chow sessions. Haloperidol (0.1 mg/kg) decreased strawberry milk consumption (a) and did not significantly change chow consumption (b). We also analyzed the data by normalizing performance to baseline (no vehicle or drug injection) sessions. After normalization, there was still a significant decrease in strawberry milk consumption (c) and also an increase in chow consumed (d). \**p* < 0.05; \*\*\**p* = 0.001. N = 15.

As with tolcapone, we also compared the effects of haloperidol to vehicle treatment as a percentage change from baseline performance. We found that haloperidol also decreased the amount of strawberry milk earned (*t_15_* = 4.657, *p* = 0.0003; Figure 3c) and it also significantly increased chow consumption (*t_15_* = 2.221, *p* = 0.0422; Figure 3d).

## Discussion

Here, we describe a series of experiments using a modified touchscreen-based ERC task. In agreement with previous studies using multiple ERC tasks, we found that male mice modulate their consumption of a highly palatable (high value) food source based on the amount of work required to attain the food. As the work requirement for the high value food increased, mice both decreased their consumption of the high value food and increased the consumption of freely available standard chow. We validated our novel ERC task by using the compound haloperidol as a positive control. We found that haloperidol decreased the amount of the high value food earned and consumed and increased the amount of chow consumed. These effects of haloperidol have been seen previously in ERC tasks and implicate our task as assaying similar neurobiological constructs as previously described ERC protocols (Salamone et al 1994, Salamone et al 1991, Yang et al 2020). Additionally, we tested the effects of the COMT inhibitor tolcapone and found that it did not significantly modulate ERC performance at any of the doses tested.

To our knowledge, the experiments we describe here are the first report of the effects of COMT inhibition alone in an ERC task. Tolcapone had been tested in a progressive ratio/concurrent chow ERC task in combination with the vesicular monoamine transporter (VMAT) inhibitor tetrabenazine and it did not modify the decrease in progressive ratio responding caused by tetrabenazine (Randall et al 2014). These data indicate that COMT activity does not regulate ERC performance to a significant degree in male mice.

It is still possible that COMT may be critically involved in ERC behavior and our experimental conditions were just not optimized to assay the effects of COMT inhibition. For example, we chose to focus on an FR4/concurrent chow ERC task and COMT inhibition may modulate performance in ERC tasks that use different reinforcement schedules or different types of effort (Randall et al 2014, Salamone et al 1994, Yohn et al 2015b). We also used an acute, single-dose treatment strategy to modulate COMT activity, so we did not address the effects of long-term alterations in COMT function. Utilization of genetic models of variable COMT activity may demonstrate a specific role for COMT in ERC performance (Yang et al., 2018). Additionally, COMT function has been shown to be sexually dimorphic in both humans and rodents (Sannino et al 2017), so it is possible COMT may significantly regulate ERC performance in females. Future studies will be designed to address potential sex differences in ERC performance.

We believe the effects of tolcapone reported here are an accurate representation of the effects in this population (young, male C57BL/6J mice) because our validation data indicate we used a robust ERC task. Our current protocol was a modified version of a previously published touchscreen ERC assay (Heath et al 2015). Also, haloperidol produced the expected decrease in preference for the high-effort high-value option seen in traditional ERC tasks (Salamone et al 1991) and a recently developed touchscreen version (Yang et al 2020).

Our data suggest an interesting dichotomy in the role of COMT in reward-seeking behavior. There are very clear effects of COMT modulation on intertemporal choice and delay discounting (Boettiger et al 2007, Kayser et al 2012) while our data show no effect on ERC behavior. There is evidence that distinct neural circuits regulate delay and effort discounting (Klein-Flugge et al 2016, Prevost et al 2010). Additional experimentation will be required to determine the exact role of COMT function in reward-seeking behavior, particularly related to effort and delay discounting. Nevertheless, to the extent that ERC behavior is critically dependent on dopamine signaling in the ventral striatum, our data are consistent with COMT being a minor player in striatal DA function.

COMT inhibition has been proposed as a therapeutic strategy for combating cognitive impairment, particularly disorders of cognitive control (Apud & Weinberger 2007). There is evidence in humans and rodents that COMT inhibition may be beneficial for specific populations (Gasparini et al 1997, Kayser et al 2012, McCane et al 2014) or on specific cognitive tasks (Tunbridge et al 2004, Byers et al 2019). Determining the utility of COMT inhibitors will require data showing the effects of these compounds across multiple cognitive domains, so that clinical populations that would benefit from this class of compounds can be identified. These experiments represent a potentially important contribution to this literature as the first report on the effects of COMT inhibition on effort-related decision-making.

## Acknowledgements

The authors thank Ingrid Buchler and Noelle White for their technical assistance.

